# p115RhoGEF activates RhoA to support tight junction maintenance and remodeling

**DOI:** 10.1101/2022.06.08.495306

**Authors:** Shahana A. Chumki, Lotte M. van den Goor, Benjamin N. Hall, Ann L. Miller

## Abstract

In vertebrates, epithelial cell-cell junctions must rapidly remodel to maintain barrier function as cells undergo dynamic shape-change events. Consequently, localized leaks sometimes arise within the tight junction (TJ) barrier, which are repaired by short-lived activations of RhoA, called “Rho flares”. However, what activates RhoA at leak sites remains unknown. Here, we asked which guanine nucleotide exchange factor (GEF) localizes to TJs to initiate Rho activity at Rho flares. We find that p115RhoGEF locally activates Rho flares at sites of barrier leaks and TJ loss. Knockdown of p115RhoGEF leads to diminished Rho flare intensity and impaired TJ remodeling. p115RhoGEF knockdown also decreases junctional active RhoA levels, thus compromising the apical actomyosin array and junctional complex. Furthermore, p115RhoGEF is necessary to preserve global TJ barrier function by promoting local leak repair and preventing repeating leaks. In all, our work demonstrates a central role for p115RhoGEF in activating junctional RhoA to preserve barrier function and direct local TJ remodeling.

## Introduction

Epithelial barrier function is critical for tissue development and homeostasis, selectively regulating the movements of molecules and ions across the barrier while preventing invasion of harmful pathogens (Marchiando *et al*., 2010). To execute effective barrier function, polarized epithelial cells must establish and maintain adhesion to one another. In vertebrates, cell-cell adhesion is regulated by the apical junctional complex, including tight junctions (TJs) and adherens junctions (AJs) (Hartsock and Nelson, 2008). Transmembrane TJ proteins partner with those in neighboring cells to form branching, strand-like networks that create a selectively permeable barrier (Varadarajan *et al*., 2019; Saito *et al*., 2021). Cytoplasmic scaffolding proteins connect TJ strands to an apical array of actomyosin, functionally linking barrier function with cell mechanics (Arnold *et al*., 2017; Van Itallie *et al*., 2017)

Dynamic cell shape change events, like cell extrusion and cell division, challenge cell adhesion and barrier integrity (Guillot and Lecuit, 2013; Gudipaty and Rosenblatt, 2017). When the connection between TJs and the cytoskeleton is compromised, barrier function is disrupted, leading to increased paracellular permeability often correlating to disease states (Zeissig *et al*., 2007; Ivanov *et al*., 2010). In response to dynamic cell shape change, TJs actively remodel to maintain their connection and barrier integrity (Rosenblatt *et al*., 2001; Higashi *et al*., 2016; Stephenson *et al*., 2019). However, the mechanisms by which TJs coordinate junction remodeling with cell shape change are not fully understood.

The small GTPase RhoA is central director of epithelial shape change (Thumkeo *et al*., 2013; Arnold *et al*., 2017; Cavanaugh *et al*., 2020). RhoA acts as a molecular switch, requiring guanine nucleotide exchange factors (GEFs) to promote RhoA’s switch from an “inactive” GDP-bound state to an “active” GTP-bound state (Rossman *et al*., 2005). Spatiotemporally precise activation of Rho to support specific cellular events is often achieved by recruitment and/or activation of specific RhoGEFs (Garcia-Mata and Burridge, 2007; Fritz and Pertz, 2016). Active RhoA stimulates contractile F-actin assembly and Myosin II filament formation, forming actomyosin contractile arrays that drive a range of cell-scale and tissue-scale contractile events.

In epithelial tissues, active RhoA localizes to apical cell-cell junctions at steady-state tension (Terry *et al*., 2011; Reyes *et al*., 2014; Priya *et al*., 2015). In high-tension environments, for example due to acute tensile stress or apoptosis, junctional active RhoA accumulates across epithelial monolayers to promote downstream signaling that preserves epithelial integrity (Acharya *et al*., 2018; Duszyc *et al*., 2021). RhoA also mediates local TJ remodeling in response to leaks in the TJ barrier associated with junction elongation, which were detected in *Xenopus laevis* embryos (Stephenson *et al*., 2019). In response to these barrier breaches, RhoA is activated in transient, local bursts, called “Rho flares” (Stephenson *et al*., 2019). Rho flares rapidly repair the TJ barrier through localized actomyosin accumulation, which promotes junction contraction and TJ protein reinforcement (Stephenson *et al*., 2019). Recent work revealed that intracellular calcium flashes, along with the mechanical stimulus of junction elongation, are early events in the Rho flare signaling pathway (Varadarajan *et al*., 2022). These calcium flashes, mediated through mechanosensitive calcium channels, are required for sustained RhoA activation to support efficient repair and reinforcement of the TJ barrier (Varadarajan *et al*., 2022). However, a crucial step of the Rho flare TJ remodeling pathway remains unclear: what activates RhoA flares?

Here, we sought to identify which GEF(s) localize to TJs and directly activate Rho flares. We first investigated the localization of four candidate RhoGEFs that either contain an rgRGS (<u>R</u>ho<u>G</u>EF <u>R</u>egulator of <u>G</u>-protein <u>S</u>ignaling) domain, or a different domain known to function similar to rgRGS domains. We find that p115RhoGEF localizes to Rho flares during local TJ remodeling. Knockdown of p115RhoGEF decreases Rho flare intensity, leading to impaired TJ remodeling. Furthermore, p115RhoGEF is also required for proper baseline levels of junctional active RhoA, which supports the maintenance of the apical actomyosin array, TJs, and AJs. Finally, we show that p115RhoGEF guards against repeating local barrier leaks and is critical for maintaining global TJ barrier function.

## Results

### p115RhoGEF localizes to Rho flares during tight junction remodeling

To determine which GEF(s) activate Rho flare-mediated TJ repair, we investigated four candidate GEFs: p115RhoGEF (also known as Arhgef1 or Lsc), LARG (Arhgef12), PDZRhoGEF (Arhgef11), and p114RhoGEF (Arhgef18) using a localization screen in gastrula-stage (Nieuwkoop and Faber stage 10-11) *Xenopus laevis* embryos. These candidates were selected because they either possess an rgRGS domain or an alternate domain that can interact with Gɑ12/3 (Fukuhara *et al*., 2001; Martin *et al*., 2016). Since *X. laevis* frogs are allotetraploid, we considered published gastrula-stage gene expression levels (Session *et al*., 2016) for each candidate GEF to determine which alloallelle (S vs. L) to investigate (**Figures 1D and S1A**). To screen candidate GEFs, embryos were injected with mRNAs encoding the probe for active RhoA (mCherry-2xrGBD (Benink and Bement, 2005; Davenport *et al*., 2016), fluorescently tagged ZO-1 to mark TJs, and the candidate GEFs C-terminally tagged with mNeon. Live imaging revealed the localization of each of the candidate GEFs relative to Rho flares. PDZRhoGEF.S and p114RhoGEF.S both localized to apical junctions but not at the sites of Rho flares, while LARG.L remained cytosolic (**Figure 1A**). Notably, we discovered that p115RhoGEF (both p115.S and p115.L) co-localized with active Rho at sites of Rho flares (**Figures 1A and S1B**). Time lapse imaging demonstrated that weak baseline p115.S signal at apical junctions increased locally at sites of ZO-1 loss (**Figures 1, B and C, S1B, and Video 1**). As the Rho flare occurs, the peak intensity of p115.S signal preceded the peak of active RhoA signal by ∼20 seconds, and both quickly returned to baseline following ZO-1 reinforcement (**Figure 1C**). Z-depth imaging revealed that p115.S localizes strongly at the apical surface, with p115.S signal partially overlapping with the TJ protein ZO-1 (**Figure S1, C-C”**). Taken together, our data indicates that p115RhoGEF localizes to Rho flares during TJ repair and reinforcement.

**Figure 1:**
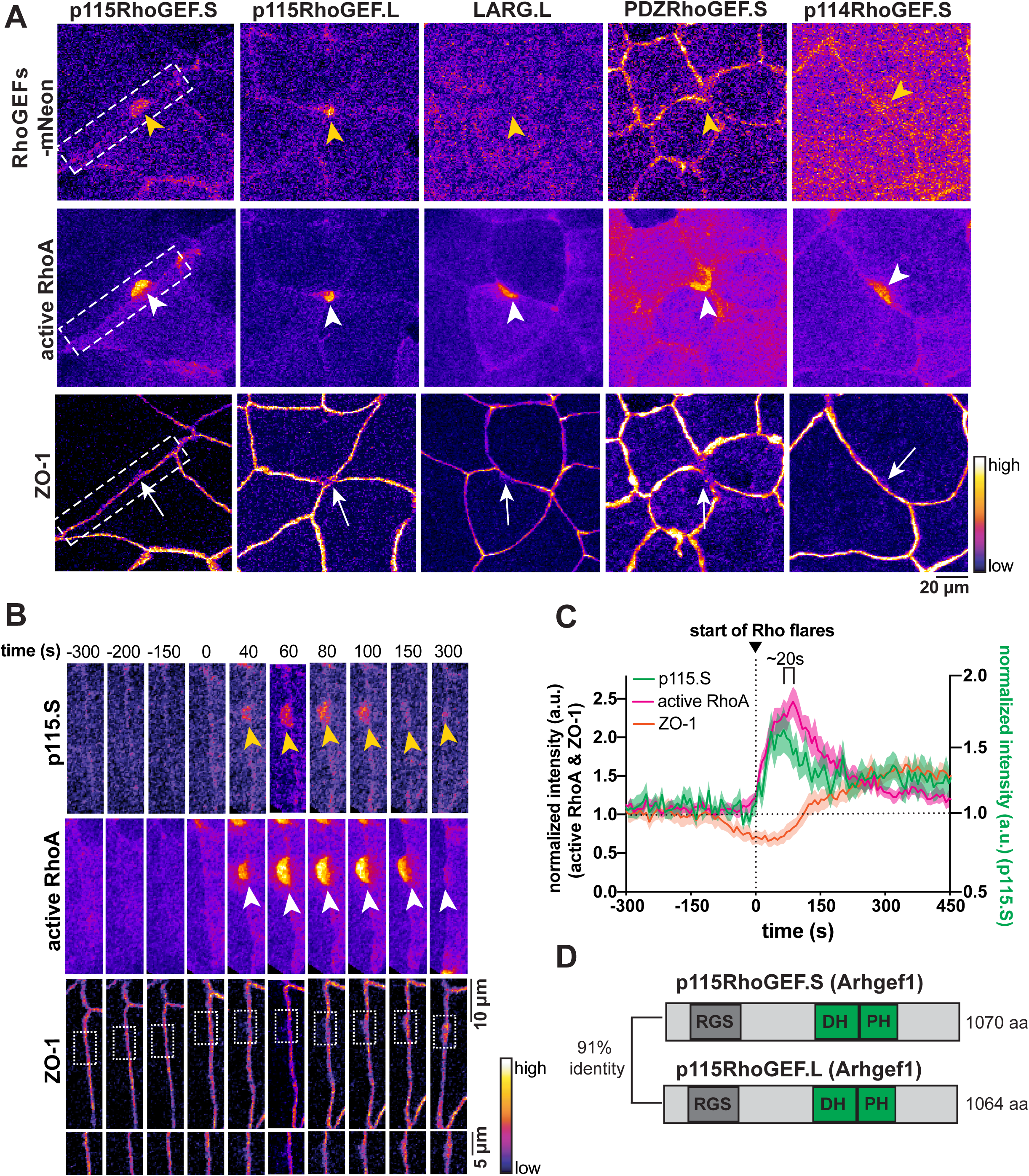
p115RhoGEF localizes to Rho flares during tight junction remodeling. (A) Cells expressing fluorescently tagged candidate RhoGEFs (p115RhoGEF.S-mNeon, p115RhoGEF.L-mNeon, LARG.L-mNeon, PDZRhoGEF.S-mNeon, and p114RhoGEF.S-mNeon, all Fire LUT), active RhoA probe (mCherry-2xrGBD, Fire LUT), and fluorescently tagged ZO-1 (BFP-ZO-1, Fire LUT). Loss of ZO-1 (white arrows) corresponds to sites of active RhoA flares (white arrowheads). Yellow arrowheads indicate sites of Rho flares in tagged candidate RhoGEF images. (B) Time lapse montage of junction indicated by the white dashed boxes in A. Local increase in p115.S (yellow arrowheads) occurs at site of ZO-1 loss (white boxed regions enlarged below). p115.S signal intensity peaks with the peak of active RhoA (white arrowheads). (C) Mean normalized intensity of p115.S, active RhoA, and ZO-1 at sites of ZO-1 loss, quantified from B and additional videos. p115.S signal intensity peaks ∼20s before the peak of active RhoA. Shading represents S.E.M. n=15 flares, 5 embryos, 4 experiments. (D) Protein domain diagram (RGS: regulator of <u>G</u>-protein <u>s</u>ignaling; DH: Dbl homology; PH: pleckstrin homology) of *Xenopus laevis* p115RhoGEF.S and p115RhoGEF.L (Arhgef1). The alloalleles (S and L) exhibit 91% amino acid identity.

### p115RhoGEF is required for Rho flares and tight junction remodeling

p115RhoGEF is known to regulate RhoA activation at cell-cell junctions in several morphogenetic contexts. For example, during cell intercalation in the *Drosophila* ectoderm, dRhoGEF2 (an ortholog of p115RhoGEF) activates Rho1 at the medial-apical cell surface (Garcia De Las Bayonas *et al*., 2019). During cell extrusion, p115RhoGEF activates RhoA at AJs to maintain cell adhesion and direct the orientation of apoptotic cell extrusion (Slattum *et al*., 2009; Duszyc *et al*., 2021). Whether p115RhoGEF is required for local Rho flare activation during TJ remodeling is unknown. To test this question, we decreased p115RhoGEF protein expression using antisense morpholinos oligos (MOs) that block translation of *Xenopus* p115RhoGEF (**Figure S2, A-D**). Initially, we aimed to knock down both p115.S and p115.L alloalleles. However, this double alloallele knockdown (p115.S+L KD) completely abolished Rho flares, preventing analysis of Rho flare frequency and behavior (**Figure S2E**). Therefore, going forward, we knocked down a single alloallele (p115.S KD), which resulted in detectable Rho flares and higher frequency of Rho flares compared to p115.S+L KD (**Figure S2F**). Rho flares in p115.S KD embryos displayed reduced intensity and duration of active RhoA at sites of TJ protein loss (**Figure 2, A-C’, and Video 2**). This reduction in active RhoA at flares led us to test whether downstream TJ repair is also compromised (Stephenson *et al*., 2019; Varadarajan *et al*., 2022). Indeed, the rapid junction contraction that immediately follows Rho flares in control embryos was absent in p115.S KD embryos (**Figure 2, D-D’**). We also measured a significant reduction in Occludin reinforcement following TJ breaks in p115.S KD embryos compared to controls (**Figure 2, B and E-E’**). This phenomenon is clearly visualized with kymographs of individual junctions, where controls exhibit robust Rho activation, junction contraction, and Occludin reinforcement, whereas p115.S KD embryos exhibit repeating Rho flares that failed to promote junction contraction or Occludin reinforcement (**Figure 2, F and G**). Of note, these clear effects occur with just a partial knockdown of p115RhoGEF levels due to the single p115.S KD. Together, these results demonstrate that p115RhoGEF is required for Rho flare activation and efficient TJ repair and remodeling.

**Figure 2:**
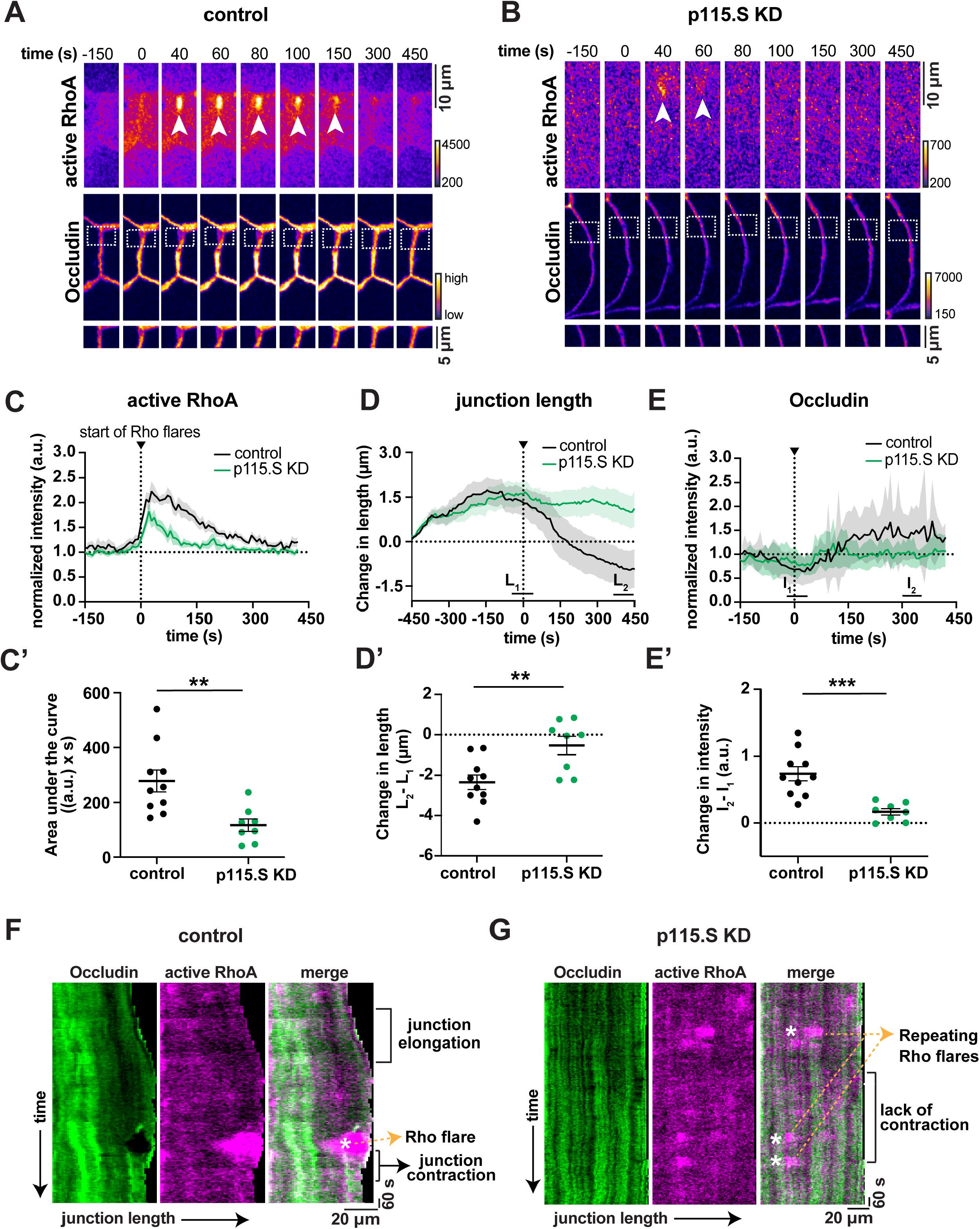
p115RhoGEF is required for Rho flares and tight junction remodeling. (A-B) Time lapse montage of active RhoA (mCherry-2xrGBD, Fire LUT) and GFP-Occludin (Fire LUT) in control embryos (A) and p115.S KD embryos (B). p115.S KD flares exhibit decreased Rho activity (white arrowheads) and Occludin reinforcement (white dashed boxes, enlarged below) in comparison to control. Time 0 represents the start of the Rho flare increase. (C-E) Graphs of mean normalized intensity of active RhoA, Occludin (E), and change in junction length (D) between control (black lines) and p115.S KD flares (green lines). Shading represents S.E.M. Control: n=10 flares, 3 embryos, 3 experiments; p115.S KD: n=8 flares, 3 embryos, 3 experiments. (C’-E’) Scatter plot of area under the curve for active RhoA (C’) was calculated from C; p=0.0047(**). Scatter plot of change in length (D’) was calculated from D. L1 and L2 represent average length of junctions from individual traces from time -20-30 s and 400-450 s, respectively; p=0.0059(**). Scatter plot of change in Occludin intensity (E’) was calculated from E. I1 and I2 represent average intensity of Occludin from individual traces from time -20-30 s and 300-350 s, respectively; p=0.0004(***). Error bars represent mean ± S.E.M.; significance was calculated using unpaired *t* tests. (F-G) Kymographs of Occludin (green), active RhoA (magenta), and merged image from representative junctions projected from vertex-to-vertex over time in control and p115.S KD embryos. Kymographs highlight repeated Rho flares, lack of Occludin reinforcement, and lack of junction contraction in p115.S KD embryos.

### p115RhoGEF is required for junctional RhoA activation to maintain the apical actomyosin array, tight junctions, and adherens junctions

Knocking down p115RhoGEF revealed another surprising phenotype: compared to controls, p115RhoGEF KD cells displayed a dramatic increase in cell size (**Figure S3A**). Therefore, we asked if the cell size changes occurred in response to dysfunctional actomyosin contraction at apical cell junctions or cytokinesis defects.

We first examined three components of junction contractility: 1) active RhoA, 2) F-actin, and 3) Myosin II. Live imaging revealed a reduction in the baseline levels of active RhoA at apical junctions in p115.S+L and p115.S KD embryos (**Figure 3, A and B**). Notably, junctional active RhoA intensity was partially rescued when p115.S KD embryos were injected with a p115.S mutant that could not be targeted by the MO (p115.S wobble) (**Figures 3, A and B, S2A**). Additionally, we found that expression of the p115.S wobble mutant partially rescued cell size defects in p115.S KD embryos (**Figure S3A**). Fixed imaging revealed significantly decreased F-actin and phospho-Myosin II (P-MLC) intensity at apical junctions in p115RhoGEF KD embryos, which was rescued by expression of p115.S wobble (**Figure 3, C-E**). Since cytoskeletal rearrangements are crucial for cell migration and intercalation during tissue remodeling (Huebner and Wallingford, 2019), we predicted that this loss of actomyosin junction contractility due to p115RhoGEF KD could cause developmental defects during morphogenesis. Indeed, p115.S+L KD embryos did not progress past gastrulation, and p115.S KD embryos exhibited severe delays in body axis elongation that were partially rescued with p115.S wobble expression (**Figure S3B**).

**Figure 3:**
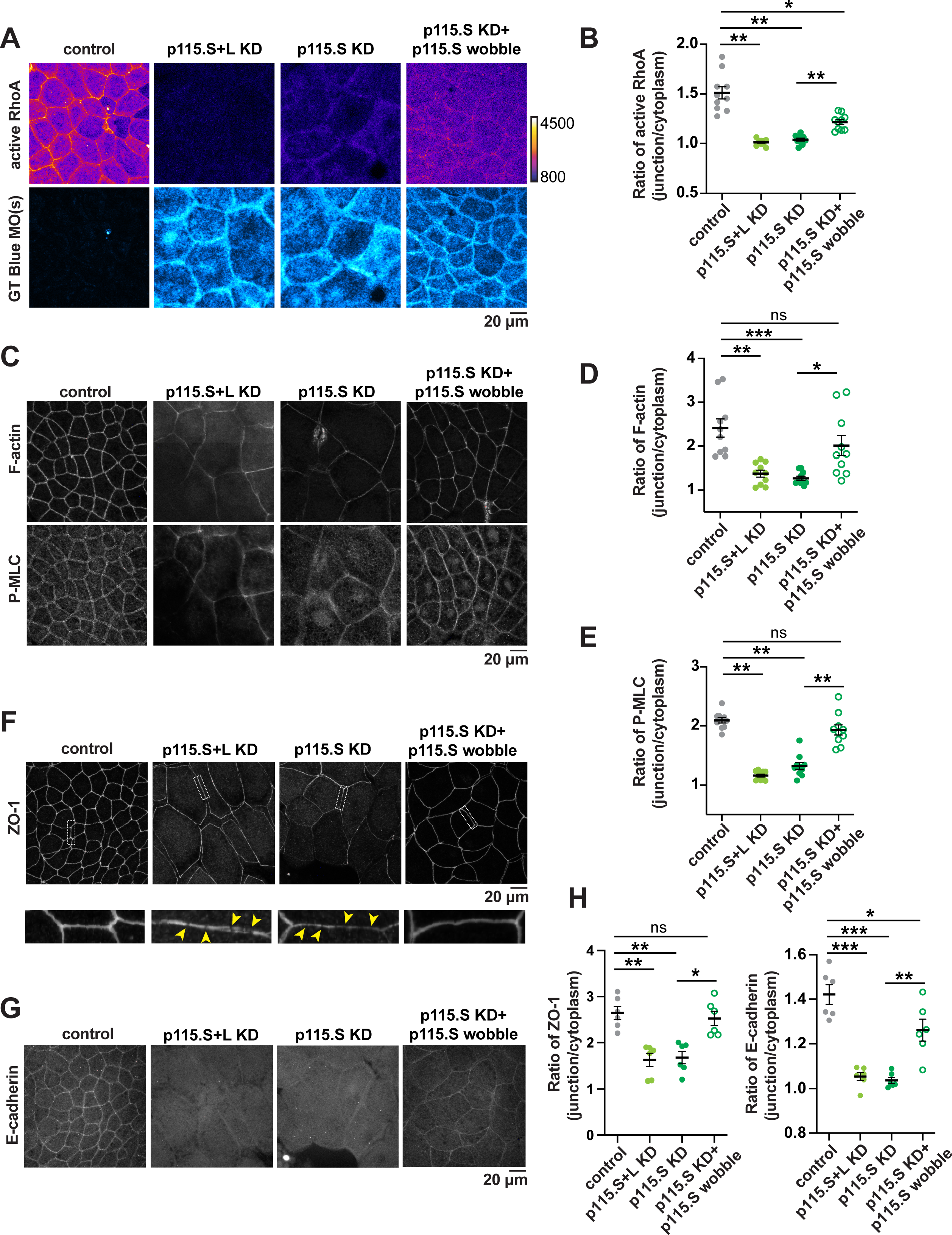
p115RhoGEF is required for junctional RhoA activation to maintain the apical actomyosin array, tight junctions, and adherens junctions. (A-B) Live imaging of cells expressing active RhoA probe (mCherry-2xrGBD, Fire LUT) in control, p115.S+L KD, p115.S KD, and p115.S KD + p115.S wobble embryos. GeneTools-Blue MO(s), when present, are shown in blue (Cyan Hot LUT). (B) Ratio of junction/cytoplasm signal of RhoA is reduced with p115RhoGEF KD. Injection of p115.S wobble partially rescues junctional active RhoA intensity. p=<0.0001(**) and p=0.003(*). Error bars represent mean ± S.E.M.; significance was calculated using unpaired *t* tests. Control: n=10 embryos, 3 experiments; p115.S+L KD: n=10 embryos, 3 experiments; p115.S KD: n=10 embryos, 3 experiments; p115.S KD + p115.S wobble: n=10 embryos, 3 experiments. (C-E) Sum projection of fixed staining of F-actin (Alexa Fluor 488 or 647 phalloidin), P-MLC (anti-Phospho-Myosin Light Chain2) in control, p115.S+L KD, p115.S KD, and p115.S KD + p115.S wobble embryos. (D) Ratio of junction/cytoplasm signal of F-actin is reduced with p115RhoGEF KD and rescued with p115.S wobble injection. p=<0.0001(***), p=0.0002(**), and p=0.0055(*). (E) Ratio of junction/cytoplasm signal of P-MLC is reduced with p115RhoGEF KD and rescued with p115.S wobble injection. p=<0.0001(**). Error bars represent mean ± S.E.M.; significance was calculated using unpaired *t* tests. Control: n=10 embryos, 3 experiments; p115.S+L KD: n=10 embryos, 3 experiments; p115.S KD: n=10 embryos, 3 experiments; p115.S KD + p115.S wobble: n=10 embryos, 3 experiments. (F-H) Sum projection of fixed staining for tight junctions (anti-ZO-1) in control, p115.S+L KD, p115.S KD, and p115.S KD + p115.S wobble embryos. White dashed rectangle of individual junctions is enlarged below; yellow arrowheads indicate discontinuities in ZO-1 signal. (G) Sum projection of fixed staining for adherens junctions (anti-E-cadherin) in control, p115.S+L KD, p115.S KD, and p115.S KD + p115.S wobble embryos. (H) Ratio of junction/cytoplasm signal of ZO-1 and E-cadherin is reduced with p115RhoGEF KD. Injection of p115.S wobble rescues ZO-1 and E-cadherin signal. For ZO-1 graph: p=0.0005(**) and p=0.0019(*). For E-cadherin graph: p=<0.0001(***), p=0.0013(**), and p=0.0349(*). Error bars represent mean ± S.E.M.; significance was calculated using unpaired *t* tests. Control: n=6 embryos, 2 experiments; p115.S+L KD: n=6 embryos, 2 experiments; p115.S KD: n=6 embryos, 2 experiments; p115.S KD + p115.S wobble: n=6 embryos, 2 experiments.

Given that disruptions in actomyosin contractility are correlated with dysfunctional junction complexes (Turner, 2000; Arnold *et al*., 2017), we next asked if TJs and AJs are compromised in p115RhoGEF KD embryos. Previous work had reported that depletion of p115RhoGEF in breast cancer epithelial cell lines leads to a reduction in the AJ protein, E-cadherin, along with enhanced cell protrusion and migration (Kher *et al*., 2014). Immunofluorescence imaging of *Xenopus* embryos demonstrated that both TJ (ZO-1) and AJ (E-cadherin) proteins were reduced at junctions under p115RhoGEF KD (**Figure 3, F-G**). Interestingly, enlarged images of ZO-1 in p115RhoGEF KD embryos reveal discontinuous signal in comparison to controls (**Figure 3F**). This effect was confirmed by live imaging of BFP-ZO-1. The ZO-1 signal in p115.S+L KD cells was patchy and discontinuous, whereas the ZO-1 signal in control cells was continuous and increased at the cell vertices (**Figure S4**). Despite careful handling of TCA-fixed embryos, we noted the presence of “cracks” between TJs in p115RhoGEF KD embryos; although we believe this is a fixation artifact, it suggests that p115RhoGEF KD embryos are more susceptible to damage (**Figure 3F**). Importantly, expression of the p115.S wobble mutant in p115.S KD embryos restores ZO-1 and partially rescues E-cadherin to levels similar to those measured in control embryos (**Figure 3H**). Likewise, p115.S wobble rescues the discontinuous ZO-1 signal and the “cracks” between TJs (**Figure 3F**).

We did not capture evidence of failed cell division, such as failed cytokinetic furrow ingression, during live imaging of p115.S KD cells (**Figure S5A**). However, live imaging with a nuclear marker did reveal larger nuclei and multinucleate cells in p115RhoGEF KD embryos when compared to controls (**Figure S5B**), so the possibility that p115RhoGEF can contribute to regulating cytokinesis in epithelial tissues should be further examined in future studies. Of note, we have previously demonstrated that perturbing MgcRacGAP, which disrupts cytokinesis (leading to larger cells), does not result in discontinuous TJ signal (Breznau *et al*., 2015). This suggests that the effect we see on TJs (**Figures 3F and H**) is specific to p115RhoGEF and is not simply due to the enlarged cell size. Collectively, these findings demonstrate that p115RhoGEF is required for junctional active RhoA to support the maintenance of the apical actomyosin array, TJs, and AJs in vertebrate epithelial tissue.

### p115RhoGEF prevents repeated local leaks to maintain global TJ barrier function

Since p115RhoGEF knockdown leads to compromised local TJ remodeling and maintenance of the apical junctional complex (**Figures 2 and 3**), we investigated if p115RhoGEF is required for epithelial barrier function. To assay barrier function, we performed a zinc-based ultrasensitive microscopic barrier assay (ZnUMBA; (Stephenson *et al*., 2019)) to detect transient TJ leaks. Briefly, this assay utilizes the fluorogenic dye FluoZin-3, which increases in intensity significantly when it encounters zinc. *Xenopus* embryos are injected with FluoZin-3 in the blastocoel cavity and mounted in their usual media plus zinc. When the barrier is intact, FluoZin-3 signal remains low and stable, but when the barrier is breached, bright FluoZin-3 signal is detected at the site of barrier damage. We found that whole-field fluorescence intensity of FluoZin-3 signal rapidly increased over time in p115RhoGEF KD embryos compared to controls (**Figure 4, A and B**), indicating a global defect in barrier function when p115RhoGEF is KD. Our previous work showed that blocking mechanosensitive calcium channels also resulted in a global barrier disruption, caused by repeated local leaks sites of TJ loss that were not efficiently repaired by the Rho flare pathway (Varadarajan *et al*., 2022). With this in mind, we examined whether local TJ leaks in p115.S KD embryos might be contributing to the global barrier defect. Kymographs of individual junctions in p115.S KD embryos demonstrated local barrier leaks, as evidenced by increased FluoZin-3 signal, that occurred repeatedly but were not followed by TJ reinforcement (**Figure 4D**). In contrast, FluoZin-3 leaks in control embryos were rapidly resolved upon junction contraction and TJ reinforcement (**Figure 4C**). Thus, our work demonstrates that p115RhoGEF helps to maintain global barrier function through activating Rho to repair local TJ leaks.

**Figure 4:**
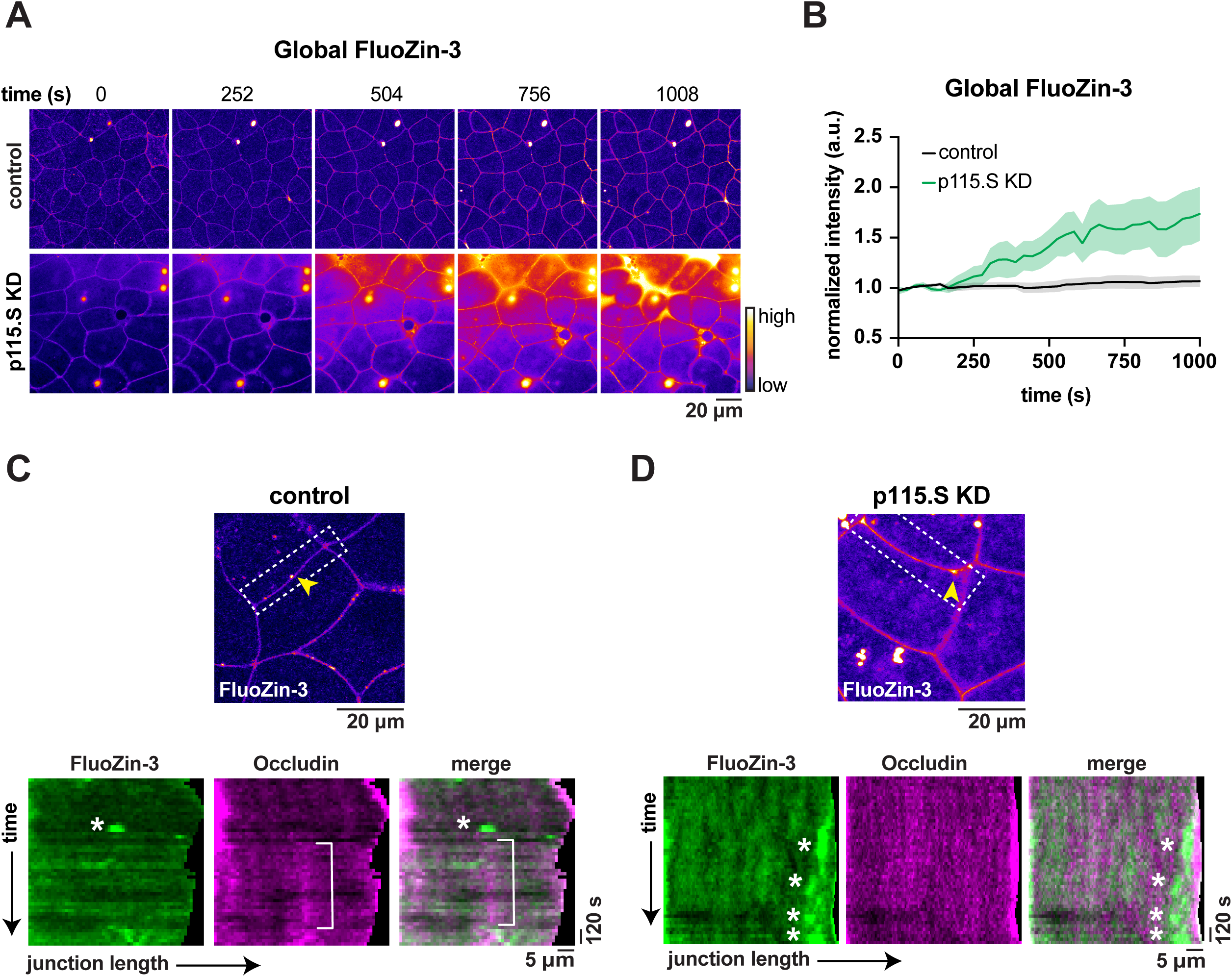
p115RhoGEF prevents repeated local leaks to maintain global TJ barrier function. (A) Time-lapse images (Fire, LUT) of global epithelial barrier permeability determined using ZnUMBA (Fluozin-3) between control and p115.S KD embryos. Time 0 represents the start of the time-lapse movie and mounting of embryos in 1mM Zinc-chloride solution. (B) Graph of mean normalized intensity of whole-field Fluozin-3 signal intensity between control (black line) and p115.S KD (green line) embryos. Global Fluozin-3 signal increases rapidly over time upon p115.S KD in comparison to control. Shading represents S.E.M. Control: n=5 embryos, 3 experiments; p115.S KD: n= 5 embryos, 3 experiments. (C-D) Cell-view of Fluozin-3 signal between control and p115.S KD embryos. White dashed rectangles represent individual junctions in kymographs. Kymographs of FluoZin-3 (green) and Occludin (mCherry-Occludin, magenta) projected from vertex-to-vertex over time in control and p115.S KD embryos. Control embryos display one Fluozin-3 leak and occludin reinforcement (white square bracket). Repeated local barrier leaks (white asterisks) and lack of Occludin reinforcement occur in p115.S KD embryos.

## Discussion

In this study, we identified *Xenopus* p115RhoGEF as a RhoGEF that activates RhoA to support TJ maintenance and remodeling (**Figure 5**). p115RhoGEF localizes to sites of TJ loss, and p115RhoGEF is required for Rho flare activation, which leads to efficient junction contraction and TJ reinforcement. Furthermore, p115RhoGEF KD compromises global junction maintenance, resulting in decreased junctional active RhoA, reduced apical actomyosin, discontinuous TJs, and reduced AJs. The functional consequence is a disruption of global barrier function brought on by repeated local leaks in the TJ. To our knowledge, the local recruitment p115RhoGEF to leak sites is the first demonstration of p115RhoGEF’s ability to regulate subcellular RhoA activation on a rapid timescale.

**Figure 5:**
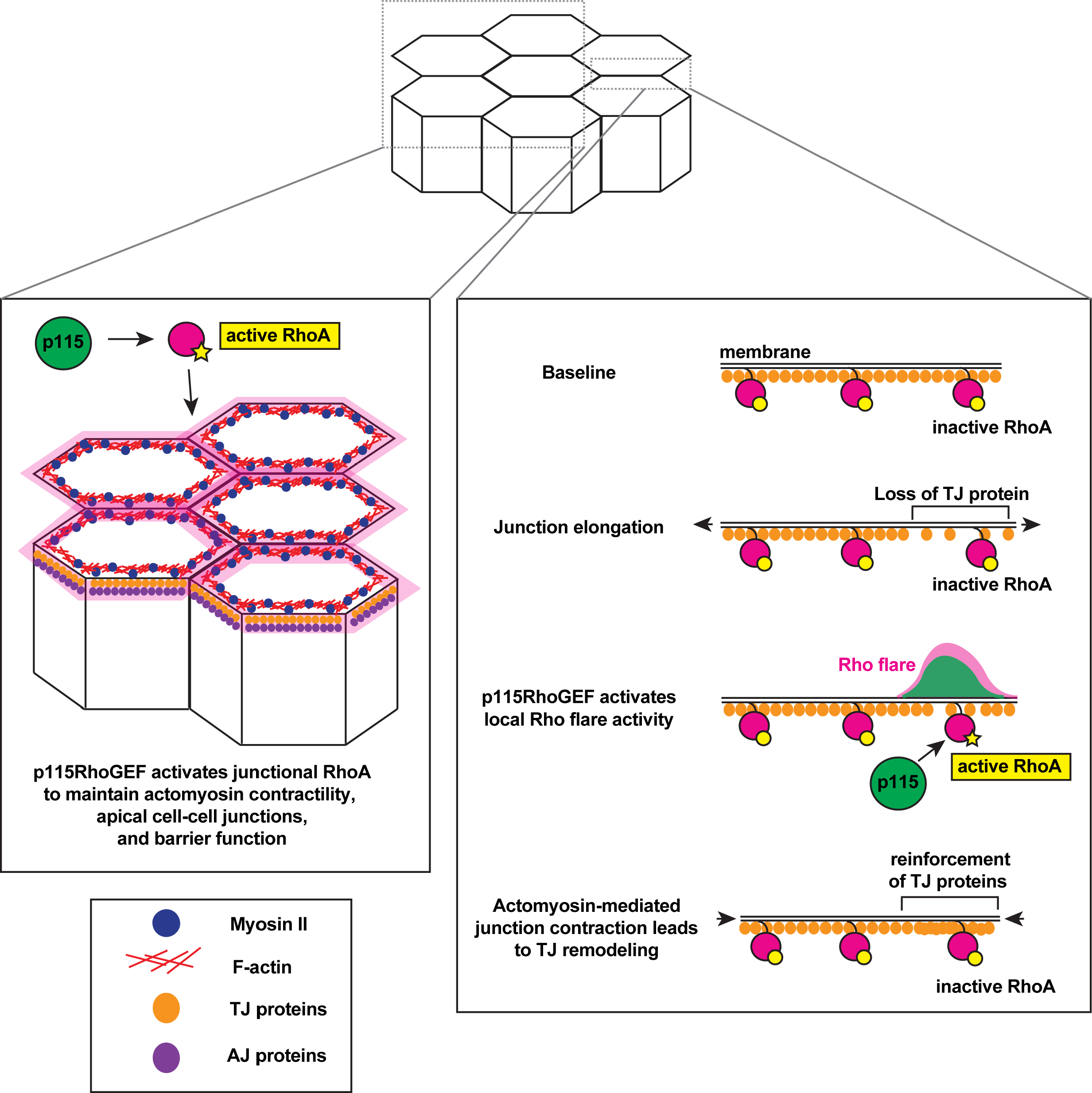
The role of p115RhoGEF in tight junction maintenance and remodeling. Model showing mechanism by which p115RhoGEF activates RhoA for TJ maintenance and remodeling. Top: 3D view of epithelial cells. Bottom left: 3D view of cell-cell junctions highlighted showing p115RhoGEF maintains baseline junctional RhoA activation to support the actomyosin array, TJs, and AJs. Bottom right: *en face* view of junction highlighted showing how p115RhoGEF recruitment and/or activation at sites of TJ damage activates Rho flares for local TJ repair and remodeling.

Given the necessity of p115RhoGEF for Rho flare activation (**Figure 2**), it is logical to consider how this GEF coordinates with other early events in Rho flare-mediated TJ remodeling, including junction elongation and cytoplasmic calcium flashes. We previously reported that the mechanical stimulus of junction elongation induces Rho flares and activates mechanosensitive calcium channel-dependent calcium influx (Varadarajan *et al*., 2022). In fibroblasts, p115RhoGEF is known to be one of the RhoGEFs, along with fellow rgRGS GEF LARG, that activates RhoA following adhesion to fibronectin (Dubash *et al*., 2007). This activation of p115RhoGEF, which happens at sites where actomyosin stress fibers insert into focal adhesions that mechanically link the cell to the extracellular matrix, may be mechanosensitive. Furthermore, in endothelial cells, tension imposed on junctional adhesion molecule A (JAM-A) activates RhoA, which leads to increased cell stiffness through kinase-mediated activation of p115RhoGEF and GEF-H1 (Scott *et al*., 2016). Thus, it follows that p115RhoGEF could be recruited and activated upon changes in local junction tension at sites of TJ damage. It will be of interest to further investigate whether Rho flare activation by p115RhoGEF occurs in a tension-dependent manner.

Another open question is how p115RhoGEF recruitment and activation coordinates with the local mechanosensitive calcium flashes that precede Rho flare activation (Varadarajan *et al*., 2022). There is evidence that protein kinase Cɑ (PKCɑ), a kinase regulated by calcium, phosphorylates p115RhoGEF, enhancing RhoA-mediated actomyosin rearrangements in endothelial cells (Holinstat *et al*., 2003). Furthermore, depletion of the calcium-sensing receptor (CaR), a member of the G protein coupled receptor (GPCR) superfamily, and/or p115RhoGEF in prostate cancer cell lines reduces aberrantly high levels of calcium-mediated choline kinase signaling, resulting in decreased tumor cell proliferation (Huang *et al*., 2011). Notably, the CaR GPCR couples to large G proteins, including Gɑ12/13. As an rgRGS protein, p115RhoGEF can be activated by Gɑ12/13, and p115RhoGEF also acts as a GTPase activating protein (GAP) toward Gɑ12/13 (Kozasa *et al*., 1998). Alternatively, there could be other tension-mediated pathways that activate p115RhoGEF by adhesion GPCRs connected to Gɑ12/13 (Storch *et al*., 2012; Vizurraga *et al*., 2020). Collectively, these studies support the notion that calcium signaling may modulate p115RhoGEF activity to ensure robust RhoA activation at sites of local barrier leaks. Future studies should investigate the molecular connection between p115RhoGEF recruitment and activation and mechanosensitive calcium signaling during the Rho flare TJ repair pathway.

In addition to the new role we report for p115RhoGEF in regulating TJ remodeling in response to local leaks in the barrier, our work also reveals that p115RhoGEF is required to maintain baseline junctional active RhoA levels to uphold the apical junctional complex and actomyosin array (**Figure 3**). Previous experiments in a breast cancer epithelial cell lines demonstrated that loss of p115RhoGEF reduces junctional E-cadherin (Kher *et al*., 2014), consistent with our results in p115RhoGEF KD *Xenopus* embryos (**Figure 3**). The previous study did not detect continuous ZO-1 signal in control cells, so it is hard to determine if p115RhoGEF depletion affected TJ organization (Kher *et al*., 2014). However, in *X. laevis* embryos, p115RhoGEF KD displayed discontinuous ZO-1 signal compared with controls as detected by both immunofluorescence and live imaging (**Figures 3 and S4**). Additionally, in this study, we provide new functional data using our sensitive live imaging barrier assay indicating that p115RhoGEF regulates barrier function by locally activating Rho to repair local barrier leaks that happen when cells change shape. It is advantageous that p115RhoGEF is recruited specifically to the site of TJ damage to locally activate Rho to reinforce TJs, as increasing Rho activity globally at apical junctions would likely break down junctions rather than repairing them.

We noted that p115RhoGEF KD cells are significantly larger than control cells (**Figure S3**), and p115RhoGEF depletion also led to increased cell size in breast cancer epithelial cell lines (Kher *et al*., 2014). Though live imaging documented p115.S KD cells in gastrula-stage *Xenopus* embryos successfully undergoing cytokinesis (**Figure S5A**), we did find evidence of multinucleate cells in p115RhoGEF KD embryos (**Figure S5B**), which were more severe in the p115.S+L KD, suggesting that loss of p115RhoGEF may lead to cytokinesis defects in epithelial tissue. p115RhoGEF has not been reported as a RhoGEF that regulates cytokinesis in isolated cells. Instead, other RhoGEFs, such as Ect2 and GEF-H1, coordinate localized RhoA activation at the cell equator by interacting with the Centralspindlin complex on the plus ends of microtubules (Ect2) or by activation via release from depolymerizing microtubules (GEF-H1) (Yuce *et al*., 2005; Birkenfeld *et al*., 2007). It would be interesting if p115RhoGEF has a role in cytokinesis that only becomes evident in tissues. In the context of a tissue, where the dividing cell needs to maintain and remodel its cell-cell junctions with its neighboring cells while pinching in two. Vinculin is recruited to sites where the contractile ring in the dividing cell is pulling on the junction with its neighboring cell to strengthen the connection of the AJ to the actin cytoskeleton (Higashi *et al*., 2016). It is possible that other proteins are also mechanosensitively recruited to these sites. Alternatively, reduced junctional actomyosin in p115RhoGEF KD cells could contribute to reduced tissue stiffness, a component that also influences cell division (Moruzzi *et al*., 2021). Future work will be needed to determine if and how p115RhoGEF contributes to successful cytokinesis in epithelial tissues.

Our study demonstrates that p115RhoGEF is important for maintaining apical cell-cell junctions (**Figure 3**). Other RhoGEFs have also been implicated in TJ formation and maintenance (Citi *et al*., 2011; Arnold *et al*., 2019). Our candidate rgRGS GEF screen revealed that like p115RhoGEF, both PDZRhoGEF and p114RhoGEF localize at apical cell-cell junctions in gastrula-stage *Xenopus* embryos; however, these two RhoGEFs were not enriched at sites of ZO-1 loss (**Figure 1**). Given that p114RhoGEF is crucial for TJ formation and protects AJs against acute tensile stress (Terry *et al*., 2011; Acharya *et al*., 2018), how does p115RhoGEF collaborate with p114RhoGEF to regulate junction formation and maintenance? One possibility is a temporal shift in which RhoGEF is the primary RhoA regulator (e.g. a handoff from p114RhoGEF to p115RhoGEF after TJ biogenesis is established). Another possibility is a spatial separation of their activities. For example, a spatially separate relationship between p115RhoGEF and p114RhoGEF orthologs is seen during morphogenesis of the *Drosophila* ectoderm, where dRhoGEF2 (ortholog of p115RhoGEF) activates Rho1 at the medial-apical surface of cells, whereas dp114RhoGEF activates Rho1 at AJs (Garcia De Las Bayonas *et al*., 2019). Another RhoGEF, GEF-H1, associates with TJs and microtubules in MDCK epithelial cells, and GEF-H1 KD reveals its necessity for selective permeability at TJs (Benais-Pont *et al*., 2003). Here, we demonstrated that p115RhoGEF regulates barrier function in *Xenopus* epithelial cells, as knockdown of p115RhoGEF compromises global barrier function and disrupts repair of local leaks (**Figure 4**). Our findings highlight the importance of p115RhoGEF in activating RhoA to promote TJ maintenance and remodeling. In future studies, it will be of high interest to continue to define the how p115RhoGEF and other RhoGEFs cooperate to regulate active Rho dynamics at cell-cell junctions.

## Materials and Methods

### DNA constructs and p115RhoGEF morpholinos

*Xenopus* rgRGS GEF sequences were synthesized by Twist Biosciences. p115RhoGEF.S (NCBI XR_001932480.2) and p115RhoGEF.L (NCBI XM_041569534.1) were ligated into the *EcoRI* and *XhoI* sites of pCS2+/C-mNeon. p115.S-wobble was generated using a gBlock from Integrated DNA Technologies and cloned into the *EcoRI* and *SphI* sites of pCS2+/p115.S-mNeon, then later ligated into the *XbaI* and *XhoI* sites of pCS2+ to make an untagged construct. LARG.L (NCBI XM_018225772.2) and PDZRhoGEF.S (NCBI XM_018233487.2) were ligated into the *BamHI* and *ClaI* sites of pCS2+/C-mNeon. p114RhoGEF.S (NCBI XM_018243648.2) was ligated into the *XhoI* and *XbaI* sites of pCS2+/C-mNeon. pCS2+/GFP-Occludin was generated by amplifying *X. laevis* Occludin from a cDNA clone purchased from Thermo Fisher (Clone ID: 7009477) and was ligated into the *BglII* and *EcoRI* sites of pCS2+/N-GFP. All constructs were verified by sequencing. The following constructs were previously described: pCS2+/mCherry-2xrGBD (probe for active Rho (Davenport *et al*., 2016)), pCS2+/BFP-ZO-1 and pCS2+/mCherry-Occludin (Stephenson *et al*., 2019), and pCS2+/mCherry-H2B (Higashi *et al*., 2016).

The p115RhoGEF antisense morpholinos (MOs) (GeneTools) were designed to target the end of the 5’UTR and start of the coding region of p115.S (MO sequence: CCAAAUUCGUCCAGAUCCAUCUC) or p115.L (MO sequence: CCAUCGUCUGAAUCCAUCUCCUG). A second p115.S MO (MO sequence: CCAAAUUCGUCCAGAUCCAUCUC) was tagged with a 3’-Gene Tools Blue fluorophore (referred to as GT BLUE MO in figures) to visualize which cells contain the MO using live or fixed imaging.

### mRNA preparation

Plasmid DNAs were linearized with *NotI*, except for BFP-ZO-1 and PDZRhoGEF.S, which were linearized with *KpnI*. *In vitro* mRNA transcription was performed from linearized plasmids using the mMessage mMachine SP6 Transcription Kit (Fisher, #AM1340) and purified using the RNeasy Mini Kit (Qiagen, #74104). RNA transcript sizes were confirmed using 1% agarose gels containing 0.05% bleach and Millennium RNA markers (Fisher, #AM7150).

### *Xenopus* embryos and microinjections

All studies conducted using *Xenopus laevis* embryos strictly adhered to the compliance standards of the U.S. Department of Health and Human Services Guide for the Care and Use of Laboratory Animals and were approved by the University of Michigan Institutional Animal Care and Use Committee. *Xenopus* eggs were collected, *in vitro* fertilized, and dejellied using methods described previously (Miller and Bement, 2009; Woolner *et al*., 2009). Embryos were stored in 0.1x Mark’s Modified Ringers (MMR) containing 10 mM NaCl, 0.2 mM KCl, 0.2 mM CaCl2, 0.1 mM MgCl2, and 0.5 mM HEPES, pH 7.4 at 15**°**C to prolong the two-cell and four-cell stages for microinjection. Embryos were microinjected as previously described (Reyes *et al*., 2014). For p115RhoGEF knockdown and replacement experiments, all four cells at the four-cell stage were first injected with either 10 nL of 2 mM p115.S and 2 mM p115.L MOs or 5 nL of 2mM p115.S MO. This was followed by a second injection at the four-cell stage of 5 nL of p115.S wobble and/or fluorescently tagged probes. Injected embryos were placed at 15**°**C for ∼24 hours until they reached gastrulation (Nieuwkoop and Faber stages 10-11). The amount of mRNA per 5 nL microinjection volume was as follows: p115.S-mNeon, 13 pg and 146 pg (for whole embryo lysate preparation); p115.L-mNeon, 13 pg; p115.S wobble-mNeon, 146 pg (1X OE, for whole embryo lysate preparation) and 292 pg (2X OE, for whole embryo lysate preparation); untagged p115.S wobble, 78 pg; LARG.L-mNeon, 12 pg; PDZRhoGEF.S-mNeon, 18 pg, p114.S-mNeon, 55 pg; mCherry-2xrGBD, 66 pg; BFP-ZO-1, 50 pg; GFP-Occludin, 8 pg; mCherry-Occludin, 8 pg; mCherry-H2B, 25 pg.

### Embryo lysates and immunoblotting

Gastrula-stage embryos (Nieuwkoop and Faber stages 10-11) were lysed as previously described (Reyes *et al*., 2014). Samples were separated on 4-20% Mini-PROTEAN TGX pre-cast gels (BioRAD, #4561094) and transferred to nitrocellulose membranes. Membranes were probed with anti-mNeon (1:100) or anti-α-tubulin (1:2500) overnight at 4**°**C in 1x Tris buffered saline (20mM Tris and 150mM NaCl) and 0.1% TWEEN-20, pH 7.6, with 5% nonfat dry milk. HRP-conjugated secondary antibodies were applied at a concentration of 1:5000. Membranes were developed using an ECL detection kit (Fisher, #32209).

### Antibodies

The primary antibodies used for western blotting were anti-mNeon (Chromotek, #32f6) and anti-α-tubulin (Sigma, T9026). Secondary antibodies used for Western blotting were horseradish peroxidase (HRP)-conjugated anti-mouse (Fisher, #PRW4021). The primary antibodies used for immunofluorescence staining were as follows: anti-Phospho Myosin Light Chain 2 (Cell Signaling Technology, #3671), anti-ZO-1 (Invitrogen, #61-7300), and anti-E-cadherin (Developmental Studies Hybridoma Bank, #5D3-C). The secondary antibodies used for immunofluorescence staining were as follows: anti-rabbit-Alexa Fluor 647 (Invitrogen, #A-21244), anti-rabbit-Alexa Fluor 568 (Invitrogen, #A-11011), and anti-mouse-Alexa Fluor 568 (Invitrogen, #A-11004).

### Immunofluorescence staining

#### F-actin (phalloidin) and P-MLC

Albino embryos were injected with control (water), 2 mM of p115.S+L MOs, 2 mM p115.S MO, or 2 mM p115.S MO and untagged p115.S wobble. Embryos were fixed at gastrula stage with paraformaldehyde (PFA) as previously described (Arnold *et al*., 2019). Embryos were fixed overnight at room temperature (RT) with a mixture of 1.5% PFA, 0.25% glutaraldehyde, 0.2% Triton X-100, and either Alexa Fluor 647 phalloidin (1:1000, Thermo Scientific, #A22287) or Alexa Fluor 488 phalloidin (1:1000, Thermo Scientific, #A12379) in 0.88x MT buffer (80 mM K-Pipes, 5 mM EGTA, and 1 mM MgCl2, adjusted to pH 6.8 with KOH). Fixed embryos were washed three times with 1x PBS, quenched for 1 hour at RT with 100 mM sodium borohydride in 1x PBS, washed again one time with 1x PBS, and bisected with a sharp scalpel to remove the vegetal hemisphere. Animal caps of the bisected embryos were blocked overnight in blocking solution (10% FBS, 5% DMSO, and 0.1% NP-40 in 1x Tris-buffered Saline). After blocking, animal caps were incubated overnight at 4°C in anti-Phospho Myosin Light Chain 2 antibody (1:100) in blocking solution, washed three times (5 minutes, 15 minutes, and then 2 hours) with blocking solution, and incubated overnight at 4°C in either anti-rabbit Alexa Fluor 568 (1:2000) and Alexa Fluor 647 phalloidin (1:1000) or anti-rabbit Alexa Fluor 647 (1:2000) and Alexa Fluor 488 phalloidin (1:1000) in blocking solution. Animal caps were washed and mounted in 1x PBS before imaging.

#### ZO-1 and E-cadherin

Albino embryos were injected with control (water), 2 mM of p115.S+L MOs, 2 mM p115.S MO, or 2 mM p115.S MO and untagged p115.S wobble. Embryos were fixed at gastrula stage with trichloroacetic acid (TCA) as previously described (Reyes *et al*., 2014). Embryos were fixed for 2 hours at RT in 2% TCA in PBS. Fixed embryos were washed three times with 1x PBS and bisected with a sharp scalpel to remove the vegetal hemisphere. Animal caps of the bisected embryos were permeabilized for 20 min at RT in 1% Triton X-100 in 1× PBS, followed by a 20 min incubation in 1× PBST (0.1% Triton X-100 in 1× PBS). Permeabilized animal caps were blocked overnight at 4°C in blocking solution (5% FBS in 1× PBST). After blocking, animal caps were incubated overnight at 4°C in anti-E-Cadherin (1:1,000) and anti-ZO-1 (1:200) antibodies in blocking solution after three washes (5 minutes, 15 minutes, and then 2 hours) in blocking solution, and incubated for 6 hours at 4°C in anti-mouse Alexa Fluor 568 (1:200) and anti-rabbit Alexa Fluor 647 (1:200) in blocking solution. Animal caps were washed and mounted in 1x PBS before imaging.

### Microscopy

#### Live imaging

Live imaging of gastrula-stage *Xenopus* embryos was performed on an inverted Olympus FluoView 1000 microscope equipped with a 60X supercorrected Plan Apo N 60XOSC objective (numerical aperture (NA) = 1.4, working distance = 0.12 mm) using mFV10-ASW software. Embryos were mounted for imaging as previously described (Reyes *et al*., 2014). Initial live imaging of mNeon-tagged candidate rgRGS GEFs, BFP-ZO-1, and mCherry-2xrGBD (active Rho probe) was captured with 8 apical Z-planes (step size 0.5 µm) acquired sequentially by line to avoid and bleed-through between channels. The scan speed was 4 µs/pixel with a 28 s time interval at a 1.5x zoom. p115.S-mNeon, BFP-ZO-1, and Rho flare movies were acquired using the 3 apical Z-planes at a 2 µs/pixel scan speed, 7 s time interval at a 1.5x zoom. p115.S MO, GFP-Occludin, and Rho flare movies were acquired with the same parameters described for p115.S-mNeon. Multi-cell view images of BFP-ZO-1 and p115.S-mNeon were captured by scanning the first 35 apical Z-planes at a 4 µs/pixel scan speed and 3.0x zoom. Mosaic images of p115.S+L MOs, p115.S MO, GFP-Occludin, and active RhoA were captured by scanning 10 apical Z-planes with a 2 µs/pixel scan speed and a 1.5x zoom. Cell view images of global active RhoA and p115RhoGEF MOs were captured by scanning the first 10 apical Z-planes at a 2 µs/pixel scan speed and 1.5x zoom. Cell view images of GFP-Occludin and mCherry-H2B were captured by scanning the first 40 apical Z-planes at a 2 µs/pixel scan speed and 1.5x zoom.

#### Fixed imaging

Bisected albino embryos were transferred into 1x PBS and mounted with the animal cap facing up within a small vacuum grease circle on a glass slide. A glass coverslip was placed over the animal cap and gently pressed down to flatten the bisected embryo. Images of embryos stained with phalloidin, P-MLC, anti-ZO-1, and anti-E-cadherin were acquired using the 30-35 Z-planes from the apical surface with a 12.5 µs/pixel scanning speed and a 1.5x zoom.

#### Live imaging barrier assay

Zinc-based Ultrasensitive Microscopic Barrier Assay (ZnUMBA) was performed as previously reported (Stephenson *et al*., 2019) with the following modifications: gastrula-stage albino embryos expressing mCherry-Occludin (control) or mCherry-Occludin and p115.S MO (p115.S KD) were microinjected into the blastocoel with 10 nL of 1 mM FluoZin-3 containing 100 µM CaCl2 and 100 µM EDTA. Embryos were allowed to recover for 1 hour at 15**°**C in 0.1x MMR. Following recovery, embryos were mounted in 0.1x MMR containing 1 mM ZnCl2 and imaged immediately using confocal microscopy using the following acquisition parameters: 8 apical Z-planes were acquired at a 4 µs/pixel scanning speed, 28 s time interval at a 1.5x zoom.

#### Images of Xenopus embryos at different developmental stages

Images of *X. laevis* embryos injected with control (water), 2 mM p115.S+L MOs, 2 mM p115.S MO, or 2 mM p115.S MO + p115.S wobble were captured with an Olympus SZX7-TR30 stereoscope using the Olympus EP50 camera and EPview app. The zoom was set to 2x for all stages except for stage 34, which required a zoom of 1.5x. Embryos were kept in a 15**°**C incubator during development and only removed for imaging.

### Image processing and quantification

Image processing and analysis were performed using Image J (FIJI). Sum intensity projections of the Z-stack were used for all quantification. Confocal images from time-lapse movies represented in the figures are sum projections of 3 apical Z-planes (1.5 μm total depth), except for multi-cell view of active RhoA (10 apical Z-planes, 5 μm total depth), multi-cell view of BFP-ZO-1 and p115.S-mNeon (35 apical Z-planes, 17.5 μm total depth), multi-cell view of GFP-Occludin and mCherry-H2B (40 apical Z-planes, 20 μm total depth), and FluoZin-3 (8 apical Z-planes, 4 μm total depth). Fixed images of phalloidin, pMLC, ZO-1, and E-cadherin are sum projections of 30 apical Z-planes (15 μm total depth).

#### Quantification of p115RhoGEF dynamics during Rho flares

Quantification of the intensity of p115RhoGEF-mNeon, BFP-ZO-1, GFP-Occludin, and active Rho probe (mCherry-2xrGBD) over time in control and knockdown experiments was performed as previously reported (Stephenson *et al*., 2019; Varadarajan *et al*., 2022) with the following modification: the small circular region of interest (ROI) had a diameter of 0.69 µm.

#### Side-view projections of ZO-1 and p115.S

Side-view projections of BFP-ZO-1 and p115.S-mNeon were acquired by capturing 35 apical Z-planes. Z-stack was opened in FIJI and was not sum projected. First, a horizontal, 0.5 µm wide segmented line was drawn perpendicular to several bi-cellular junctions (see white dashed line in Figure S1C). The 3D projection tool in FIJI was then used to generate a side-view image. Second, a rectangular ROI was drawn encompassing an individual bicellular junction, not including the vertices (see yellow dashed box in Figure S1C). The 3D projection tool was used to generate a side view along the plane of the membrane.

#### Area under the curve for Rho flares

Area under the curve for active Rho flares was quantified using GraphPad Prism 9.0 by applying area under the curve (AUC) analysis to the individual Rho flare traces. AUC analysis was applied to all points in the peaks above the baseline of 1; peaks less than 10% of the increase from the baseline to the maximum Y-value were excluded. Individual AUC values for each Rho flare for control and p115RhoGEF knockdown were plotted in a scatter plot.

#### Junction length measurements

Junction length was quantified using Occludin as a junctional marker. Using FIJI, a 0.3 μm wide segmented line was drawn on junctions with Rho flares from vertex-to-vertex every 10 frames and saved into the ROI manager. The Time Lapse and Line of Interest (LOI) interpolator plug-in then generated an ROI for each frame of the time lapse movie, accommodating for cell shape changes. Each junction was measured in triplicate and change in length was calculated by subtracting the individual values from the average of the first 65 seconds.

#### Construction of kymographs

Kymographs from vertex-to-vertex of a cell-cell junction were constructed using Occludin as a junctional marker. In FIJI, a 0.5 μm wide segmented line was drawn on junctions every 10 frames and saved into the ROI manager (a total of 200 frames for **Figure 2** and 50 frames for **Figure 4**). The Time Lapse and line of interest (LOI) interpolator plug-in then generated an ROI for each frame of the time lapse movie, accommodating for cell shape changes, and created a kymograph of those ROIs over time.

#### Frequency of Rho flares

Rho flare frequency was measured manually using FIJI, by counting the number of Rho flare occurrences at cell-cell junctions in the whole field of imaging from the start of the time-lapse movie to the end (a total of 15 minutes for each movie). Frequency was reported in graphs as flares/minute. Rho flares where the local active RhoA intensity increased and was sustained at the junctions for a span of ∼80-100s were included in the analysis.

#### Apical cell surface area measurements

Apical cell surface area was measured using GFP-Occludin. For all images, we used sum projections of 10 apical Z-slices. Some images did not have sufficient signal for automated quantification; in those cases, cells were outlined by hand using the polygon selection tool in FIJI (4 of 20 images). For all other images, cell surface area was quantified as follows using FIJI. The image was duplicated to create two copies (Image A and B). A Gaussian Blur with a radius of 15 was applied to Image B. Using the image calculator, Image B was subtracted from Image A to create Image C. A Gaussian Blur with a radius of 3 was applied to Image C. Then, thresholding was applied to Image C by manually adjusting the minimum and maximum cutoffs to maximize junction continuity. The signal was then dilated until all junctions were continuous. Then Image C was skeletonized, the signal was dilated one time, and the image was inverted. The area was measured using Analyze Particles where the size was limited from 20 -infinity microns, and cells along the edge of the image were excluded.

#### Intensity of F-actin (phalloidin), P-MLC, ZO-1, and E-cadherin

The junction/cytoplasm intensity ratios for F-actin (phalloidin), P-MLC, ZO-1, and E-cadherin were quantified in FIJI. Junctional intensity was measured with a 0.3 µm wide segmented line along bicellular junctions (excluding vertices), and cytoplasmic intensity was measured using a 10.14 µm circular ROI. For each image, we measured 5 junctions, which were averaged constituting one data point. Ratio of junctional/cytoplasmic intensity of F-actin (phalloidin), ,P-MLC, ZO-1, and E-cadherin was calculated by dividing the average intensity at the junction by the average intensity in the cytoplasm for each image.

### Global barrier function

Global barrier function was quantified as previously reported (Varadarajan *et al*., 2022) with the following modification: the baseline was normalized to 1 by dividing the individual value by the average of the first 83 seconds.

### Statistical Analysis

Unpaired Student’s *t* tests were used to determine the statistical significance of each group pairwise between control and p115.S+L KD, p115.S KD, or p115.S KD + p115.S wobble. Statistics were calculated in GraphPad Prism 9.0 software.

## Abbreviations

TJ: tight junctions
AJ: adherens junctions
GEF: guanine nucleotide exchange factor
GBD: GTPase-binding domain
rgRGS: RhoGEF regulator of G-protein signaling
MO: antisense morpholino oligos
P-MLC: Phospho-Myosin Light Chain

## Acknowledgements

We would like to thank all current and former members of the A.L. Miller laboratory for providing helpful discussion and feedback on the manuscript, especially Dr. Jennifer Landino for providing experimental advice and edits on the manuscript and Dr. Rachel Stephenson for advice on Rho flare analysis, ZnUMBA, and kymograph construction. We thank Akash Rai for assistance with cloning and initial imaging of p115.S-mNeon. This work was supported by an NIH grant (2R01 GM112794) to A.L. Miller, an NSF Graduate Research Fellowship to S.A. Chumki, and a predoctoral fellowship from the American Heart Association (906189) to L.M. van den Goor.

The authors declare no competing financial interests.

## Author contributions

S.A. Chumki and A.L. Miller conceptualized the study; S.A. Chumki, L.M. van den Goor and A.L. Miller developed the methodology; S.A. Chumki performed the majority of experiments and data analysis; L.M. van den Goor performed experiments for Figure 3 (A, C, F, G), and analyzed data for Figure S3B; B.N. Hall performed experiments for Figure S1B and investigated literature about candidate GEFs; S.A. Chumki, L.M. van den Goor, and A.L. Miller wrote the original draft of the manuscript; All authors revised the manuscript; A.L. Miller acquired funding; A.L. Miller supervised the study.

## Supplemental Figure Legends

**Figure S1:**
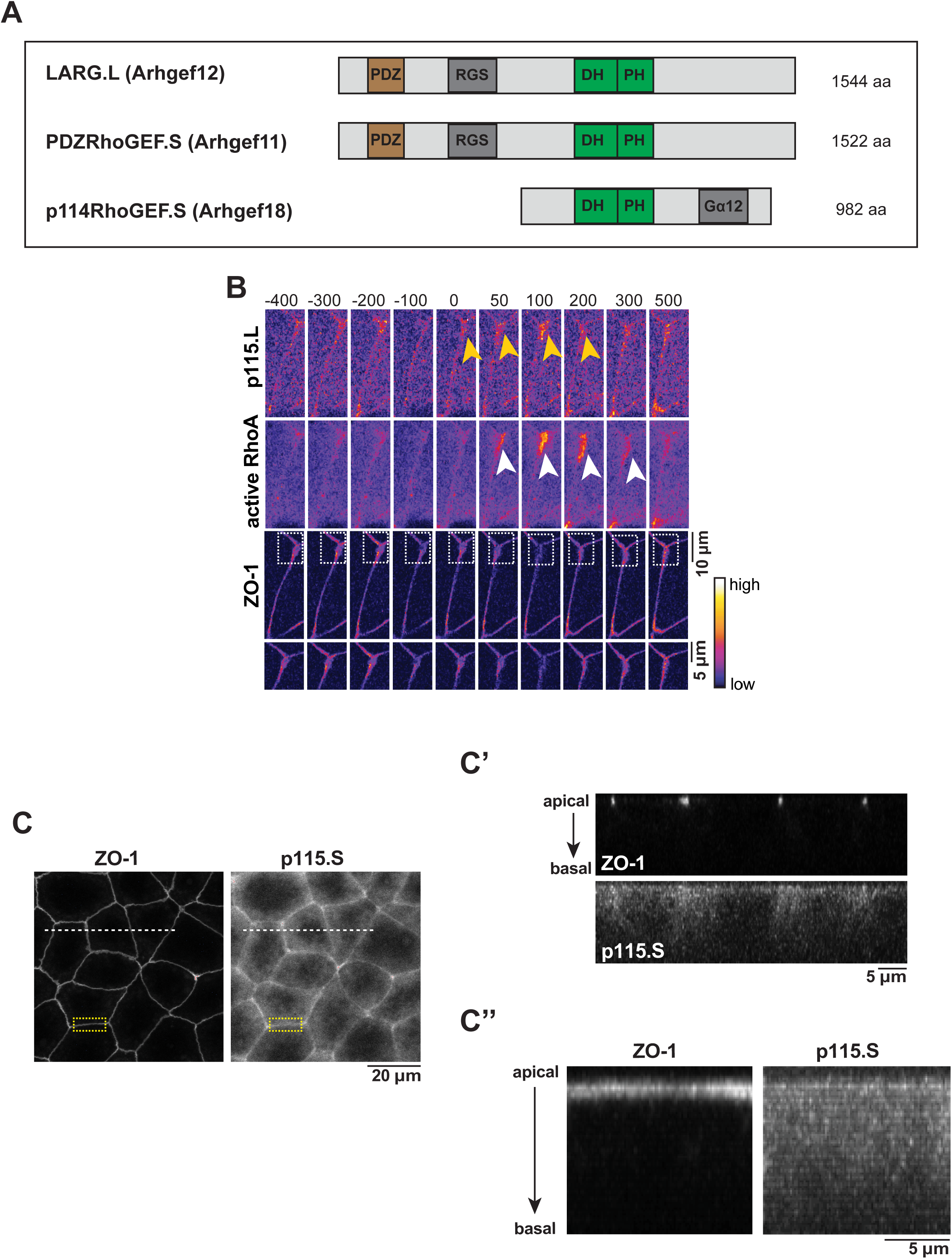
Additional characterization of candidate rgRGS GEFs. (A) Protein domain diagram (RGS: regulator of <u>G</u>-protein <u>s</u>ignaling; DH: Dbl homology; PH: pleckstrin homology; Gα12: alternative domain that binds to Gα12 of p114RhoGEF.S) of *Xenopus laevis* rgRGS GEFs: LARG.L (Arhgef12), PDZRhoGEF.S (Arhgef11), and p114RhoGEF.S (Arhgef18). (B) Time lapse montage of p115RhoGEF.L (p115.L-mNeon, Fire LUT), active RhoA probe (mCherry-2xrGBD, Fire LUT), and ZO-1 (BFP-ZO-1, Fire LUT). Local increase in p115.L (yellow arrowheads) localizes with active RhoA flares (white arrowheads) and at sites of ZO-1 loss (white boxed region is enlarged below). (C-C”) Cells expressing BFP-ZO-1 and p115.S-mNeon (both shown in grayscale). (C’) Cross-sectional side-view projection of multiple bi-cellular junctions expressing ZO-1 and p115.S (white dashed line). (C”) side-view projection of a single junction expressing ZO-1 and p115.S (yellow dashed box).

**Figure S2:**
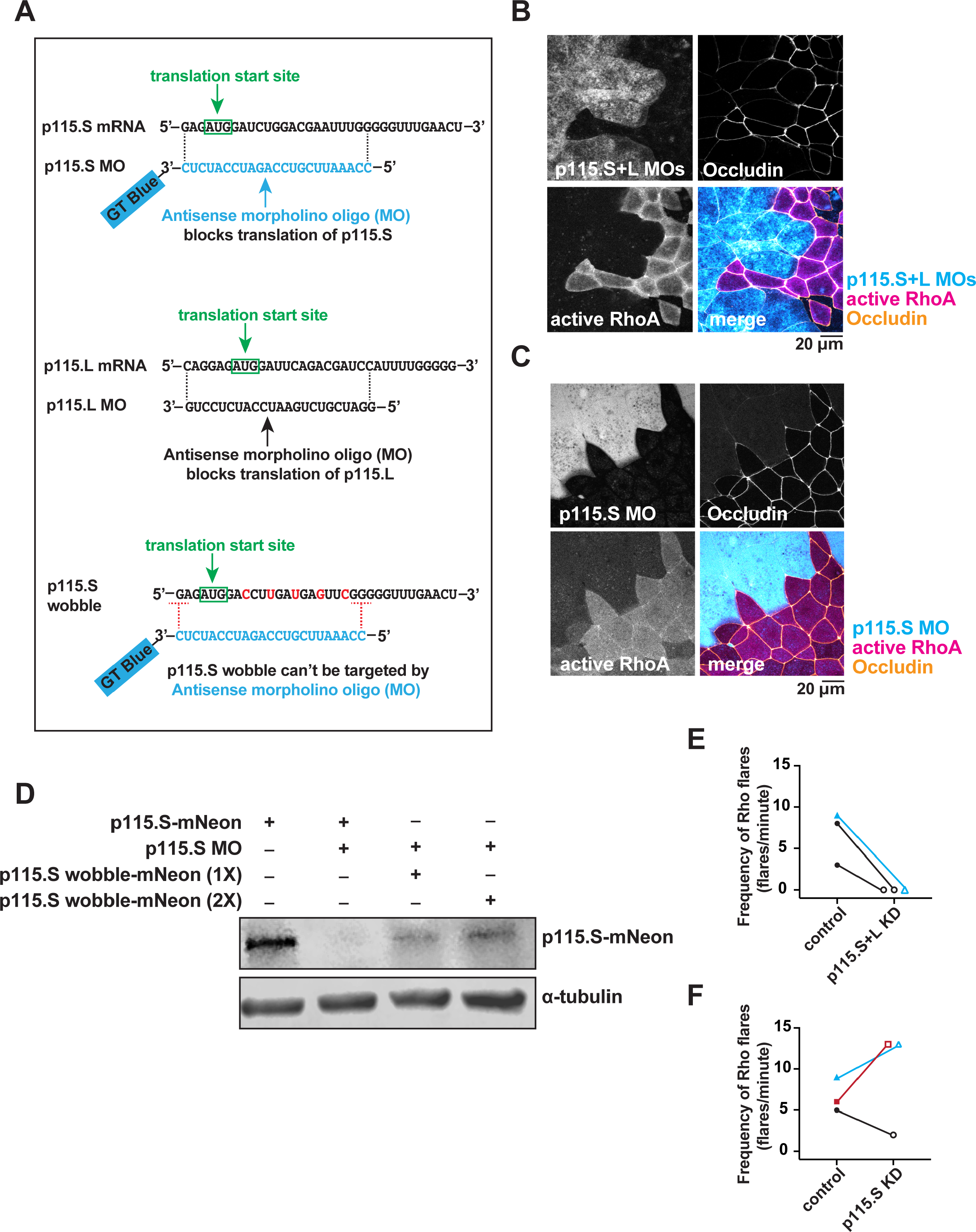
Validation of p115RhoGEF knockdown. (A) Top: schematic of 3’ Gene Tools Blue (GT Blue) tagged p115.S antisense morpholino oligo (p115.S MO) targeting p115.S mRNA to block protein translation. Middle: schematic of p115.L antisense morpholino oligo (p115.L MO) targeting p115.L mRNA to block protein translation. Bottom: Schematic of p115.S wobble mRNA containing five wobble nucleotide mutations in the protein coding region. These mutations prevent p115.S wobble from being targeted by the p115.S MO but will not affect the amino acid sequence when translated. (B) Cells mosaically expressing GeneTools-Blue p115.S MO+p115.L MO (cyan), Occludin (GFP-Occludin, orange), and active RhoA probe (mCherry-2xrGBD, magenta), as well as a merge of all three channels. (C) Cells mosaically expressing GeneTools-Blue p115.S MO (cyan), Occludin (GFP-Occludin, orange), and active RhoA probe (mCherry-2xrGBD, magenta), as well as a merge of all three channels. (D) Western blot showing p115.S-mNeon protein levels (detected with anti-mNeon antibody) when p115.S-mNeon is knocked down by injection of p115.S MO or knocked down and p115.S wobble-mNeon is expressed (1x and 2x overexpression (OE), see Materials and Methods for mRNA amounts injected). (E) Frequency of Rho flares in control and p115.S+L KD embryos. Paired experiments are matched for color and shape. Control (solid shapes): n=3 embryos, 2 experiments; p115.S+L KD (open shapes): n=3 embryos, 2 experiments. (F) Frequency of Rho flares in control and p115.S KD embryos. Paired experiments are matched for color and shape. Control (solid shapes): n=3 embryos, 3 experiments; p115.S KD (open shapes): n=3 embryos, 3 experiments.

**Figure S3:**
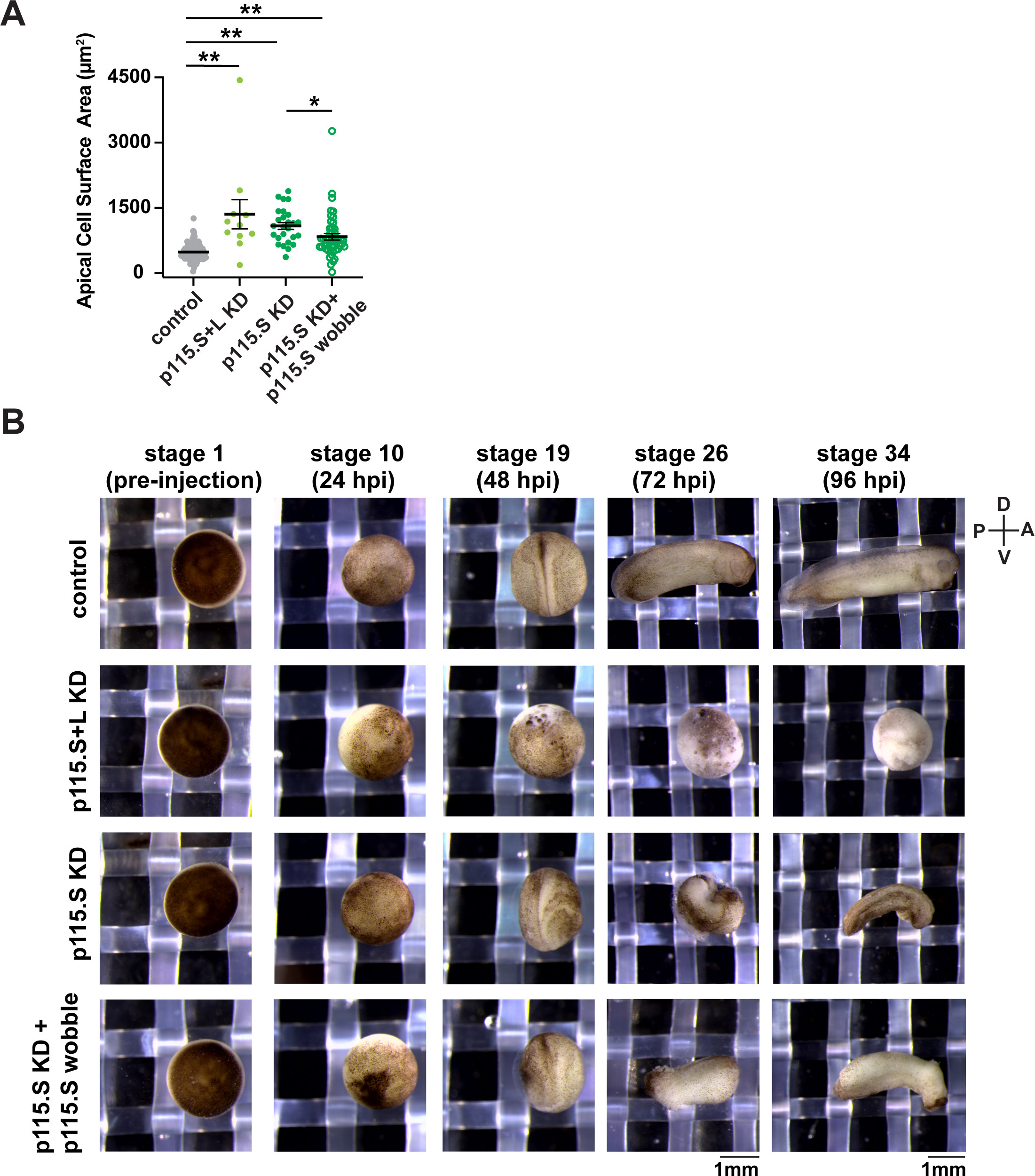
Morphological and developmental phenotypes of control and p115RhoGEF KD embryos and embryos expressing p115.S wobble. (A) Apical cell surface area quantification for control, p115.S+L KD, p115.S KD, and p115.S KD + p115.S wobble embryos. p=<0.0001(**) and p=0.0343(*). Error bars represent mean ± S.E.M.; significance was calculated using unpaired *t* tests. Control: n=143 cells, 6 embryos, 3 experiments; p115.S+L KD: n=11 cells, 3 embryos, 3 experiments; p115.S KD: n=26 cells, 6 embryos, 3 experiments; p115.S KD + p115.S wobble: n= 48 cells, 6 embryos, 3 experiments. (B) Stereoscope images of *X. laevis* embryos pre-injection, 24-, 48-, 72-, and 96-hours post injection (hpi) when kept in a 15**°**C incubator. Control, p115.S+L KD, p115.S KD, and p115.S KD + p115.S wobble embryos are shown. Approximate Nieuwkoop and Faber stages are listed as well. D=dorsal, V=ventral, P=posterior, A=anterior.

**Figure S4:**
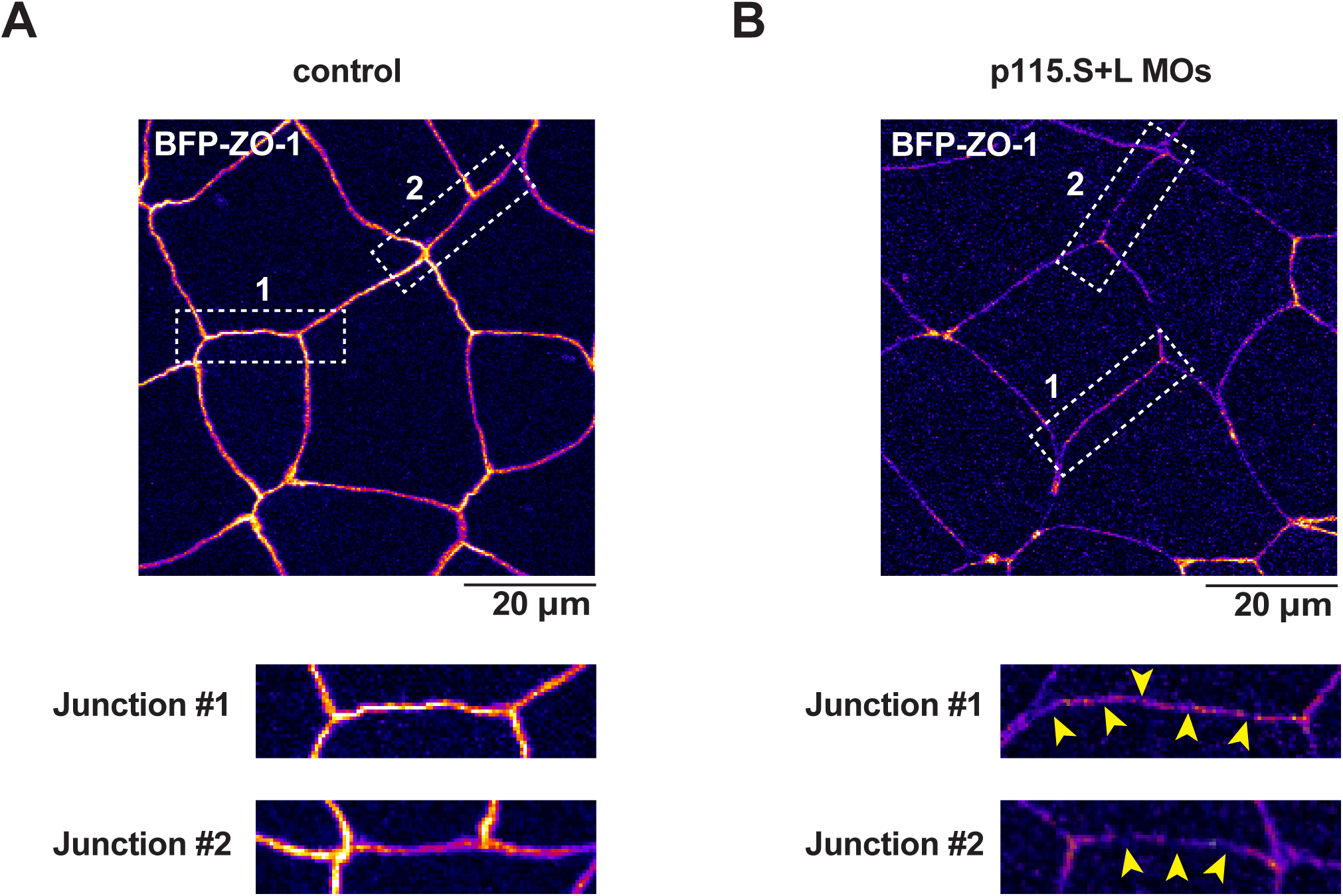
ZO-1 signal is discontinuous in live imaging of p115RhoGEF KD embryos. (A-B) Cell views of BFP-ZO-1 in control and p115.S+L KD embryos (note that 1 mM of each MO was used instead of the usual 2 mM for this experiment). White dashed rectangles are enlarged below and reveal discontinuous ZO-1 in p115.S+L KD embryos.

**Figure S5:**
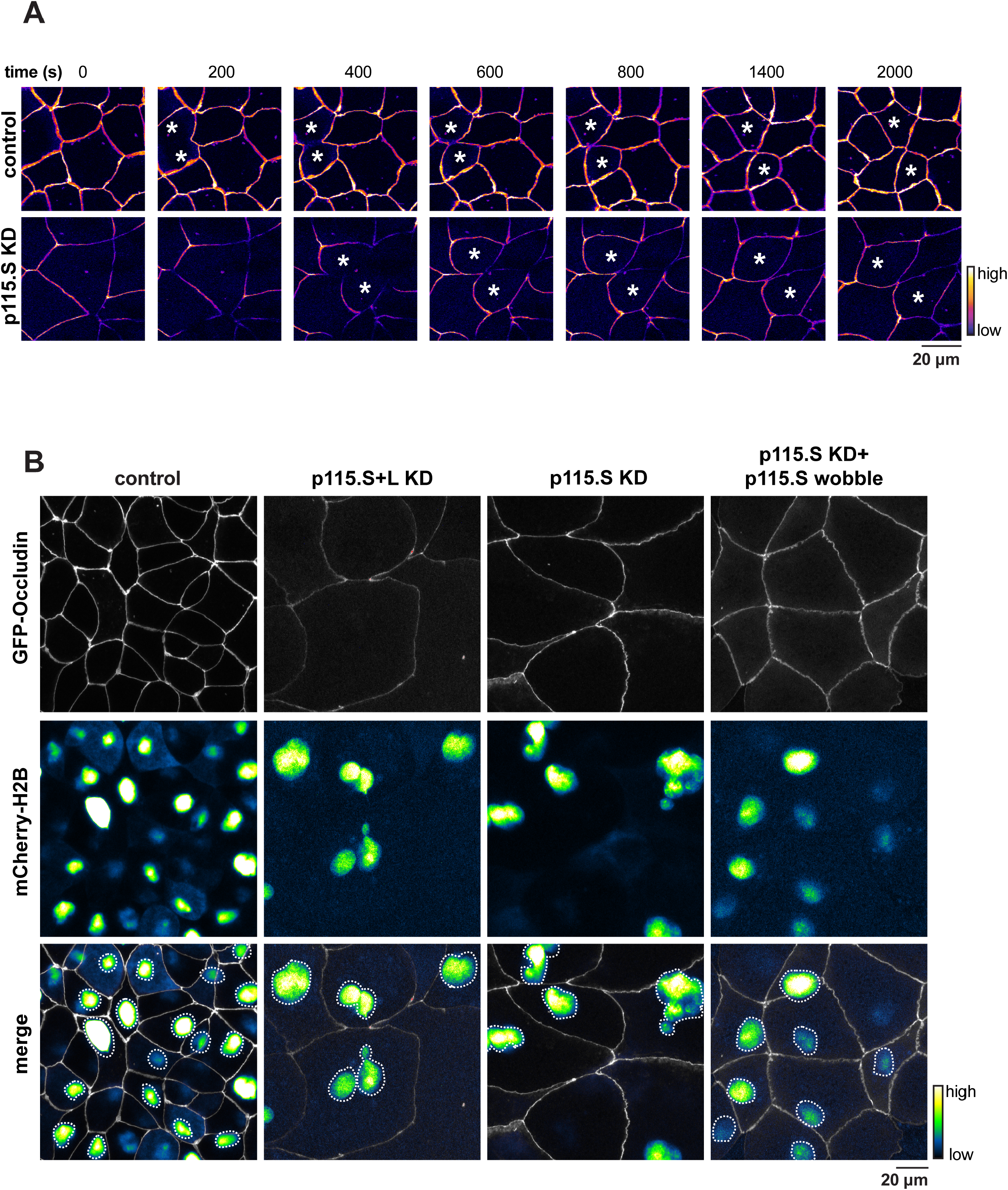
Potential effects of p115RhoGEF KD on cell division. (A) Live imaging of Occludin (GFP-Occludin, Fire LUT) in control and p115.S KD embryos. Montages show p115RhoGEF.S KD does not obviously interfere with cytokinetic furrow ingression. White asterisks indicate dividing cells. Time 0 s represents the start of furrow ingression. (B) Live imaging of GFP-Occludin (gray) and mCherry-H2B (Green Fire Blue LUT) in control, p115.S+L KD, p115.S KD, and p115.S KD + p115.S wobble embryos. White dashed lines outline nuclei.

## Supplemental Video Legends

**Video 1:** Time-lapse confocal imaging of gastrula-stage *Xenopus laevis* embryo showing that both local p115.S (p115.S-mNeon, green channel, white outlined arrows) and active Rho (mCherry-2xrGBD, magenta channel, gold arrows) increase at the site of TJ breaks (BFP-ZO-1, Fire LUT channel, white arrows). Time interval, 8 s; video frame rate, 20 fps. Related to Figure 1**, B and C**.

**Video 2:** Time-lapse confocal imaging of control and p115.S KD gastrula-stage *Xenopus laevis* embryos. Compared to control, p115.S KD movie shows repeating activation of Rho flares (mCherry-2xrGBD, magenta channel, gold arrows) along the junction. These Rho flares have reduced intensity and are followed by reduced Occludin reinforcement (GFP-Occludin, green channel, white arrows). Time interval, 7 s; video frame rate, 20 fps. Related to Figure 2**, A and B**.

